# Human cytomegalovirus nuclear capsid motility is non-directed and independent of nuclear actin bundles

**DOI:** 10.1101/641266

**Authors:** Felix Flomm, Eva Maria Borst, Thomas Günther, Rudolph Reimer, Laura de Vries, Carola Schneider, Adam Grundhoff, Kay Grünewald, Martin Messerle, Jens Bern-hard Bosse

## Abstract

Herpesvirus genome replication, capsid assembly and packaging take place in the host cell nucleus. Matured capsids leave the nucleus through a unique envelopment-de-envelopment process at the nuclear membranes called nuclear egress. How assembled and DNA-containing herpesvirus capsids reach the sites of nuclear egress is however still controversially discussed, as host chromatin that marginalizes during infection might constitute a major barrier. For alphaherpesviruses, previous work has suggested that nuclear capsids use active transport mediated by nuclear filamentous actin (F-actin). However, direct evidence for nuclear capsid motility on nuclear F-actin was missing. Our subsequent work did not detect nuclear F-actin associated with motile capsids, but instead found evidence for chromatin remodeling to facilitate passive capsid diffusion. A recent report described that human cytomegalovirus, a betaherpesvirus, induces nuclear F-actin and that the motor protein myosin V localizes to these structures. Direct evidence of capsid recruitment to these structures and motility on them was however missing. In this study, we tested the functional role of HCMV-induced, nuclear actin assemblies for capsid transport. We did not observe transport events along nuclear F-actin. Instead, reproduction of nuclear F-actin was only possible using strong overexpression of the fluorescent marker LifeAct-mCherry-NLS. Also, two alternative fluorescent F-actin markers did not detect F-actin in HCMV-infected cells. Furthermore, single particle tracking of nuclear HCMV capsids showed no indication for active transport, which is in line with previous work on alphaherpesviruses.

## Importance

Although human cytomegalovirus hardly causes disease in healthy individuals, it constitutes a major hazard to immunocompromised risk groups. Human Cytomegalovirus infests nearly all organs and can cause severe disease such as pneumonitis, colitis, encephalitis and retinitis, and can lead to serious impairments in neonates. Currently available treatments target only two steps during the viral ‘life cycle’, which makes the occurrence of viral resistance a major problem. To identify targets for pharmaceuticals, in-depth knowledge of the molecular mechanisms of the viral infection is paramount. Since the virus relies on the ability to release infectious particles from a host cell to infect another cell, its ability to translocate these particles within a cell is critical to complete the viral ‘life cycle’. This work indicates that remodeling of cellular chromatin, rather than molecular motors, enables capsid access to the nuclear membrane. Understanding the mechanism of chromatin remodeling might help in designing effective inhibitors.

## Introduction

HCMV, like other herpesviruses, creates infectious particles using a series of complex morphogenesis steps that lead to the fully assembled and infectious virion consisting of capsid, tegument, and envelope. These layers are subsequently added while the forming particles pass through several host-cell organelles (1, 2). Morphogenesis starts in the nucleus, where DNA replication, the formation of nucleocapsids, as well as the packaging of viral genomes into the latter takes place. Next, herpesvirus capsids must reach the nuclear envelope for primary envelopment and egress (3). How nuclear capsids cross the nucleoplasm is controversially discussed in the field. Earlier work suggested that herpes simplex virus type 1 (HSV-1) capsids exhibit directed nuclear motility as determined by single particle tracking. The observed directed motility could be antagonized by a putative myosin inhibitor as well as the actin-depolymerizing drug Latrunculin A, suggesting that capsids might use myosins to move actively on nuclear F-actin (4). One year later a study was able to detect F-actin in fixed rodent neuronal cells infected with pseudorabies virus (PRV), using both fluorescence microscopy as well as serial-block-face scanning electron microscopy (SBFSEM) (5). However, we recently did not detect any nuclear F-actin in fibroblasts infected with the alphaherpesviruses HSV-1, PRV, the betaherpesvirus murine cytomegalovirus (MCMV) or the gammaherpesvirus murine herpesvirus-68 (MHV-68), while nuclear capsid motility in these cells was detectable (6)

Moreover, we found that Latrunculin A was able to induce aberrant actin assemblies that seemed to unspecifically bind viral capsids and block their movement (6, 7). Since direct evidence of capsid motility along nuclear actin filaments was missing, and advances in camera technology now allow much more precise measurements, we re-evaluated previous findings. Using a custom microscope design, we acquired several thousand alphaherpesvirus nuclear capsid tracks. Analysis of these tracks showed no indication of bulk directed motility of nuclear herpesvirus capsids. Instead, we found that infection-induced chromatin remodeling allowed capsids to cross the nucleoplasm by diffusion to reach the nuclear envelope.

These findings are supported by two recent reports in which the authors were able to resolve interchromatin channels that bridge through the marginalized chromatin in HSV-1 infected cells to supposed egress sites at the nuclear envelope (8). Computational modeling using our experimentally determined diffusion coefficients indicates that these channels allow herpes-virus capsids to reach the nuclear envelope by diffusion with(9).

In discordance with these findings, a recent report showed large nuclear actin filaments in human foreskin fibroblast (HFF) cells stably expressing LifeAct-GFP-NLS and infected with HCMV (10). In addition, the authors found that prolonged incubation of infected cells with very high concentrations of the F-actin-depolymerizing drug Latrunculin A led to a defect in infectious virus production as well as their translocation to the cytoplasm, and suggested that these filaments are involved in movement of capsids to the nuclear periphery for nuclear egress. Moreover, myosin Va was implicated in nuclear egress, as the authors found a colocalization of myosin Va with the major capsid protein of HCMV at the rim of the viral replication compartment, as well as an antagonizable effect of myosin Va on nuclear capsid localization to the nuclear envelope (11).

These findings might argue for a role of nuclear F-actin in the trafficking of betaherpesvirus nucleocapsids to nuclear egress sites. However, direct evidence for active motility of capsids along nuclear filaments is missing. Recently, we developed a UL77-mGFP-tagged HCMV mutant that produces fluorescent nuclear capsids (12). We, therefore, set out to analyze the motility of HCMV nuclear capsids in relation to nuclear actin filaments by single particle tracking. While aiming at reproducing the induction of nuclear F-actin in HCMV-infected cells expressing LifeAct-NLS, we found that nuclear filament induction is dependent on the expression level and cellular localization of the utilized actin live-cell marker LifeAct. In our hands, only cells with very high expression levels of LifeAct-mCherry-NLS presented nuclear filaments, and reducing the expression levels by using a weaker promotor or utilizing a weakly expressing cell population almost completely abrogated nuclear filament occurrence. Two alternative fluorescent F-actin markers were unable to detect F-actin in HCMV-infected cells. Using electron microscopy, we could only detect nuclear F-actin in infected cells expressing mCherry-LifeAct-NLS, while in the absence of mCherry-LifeAct-NLS infected cells did not show any. Finally, deleting the NLS abolished nuclear F-actin formation, which led us to conclude that nuclear actin induction in this system is an artifact of LifeAct-NLS overexpression. In accordance, we did not find transport events along nuclear F-actin employing single particle tracking. Instead, nuclear HCMV capsids showed diffusive motility with no indication for active transport, which fits our measurements that HCMV infection also remodels the nuclear chro-matin structure to facilitate particle diffusion as described previously for members of the alphaherpesviruses. We, therefore, conclude that LifeAct-NLS serves as both an expression-level-dependent inducer and detector of nuclear actin filaments in HCMV-infected cells, while HCMV infection itself does not induce nuclear actin in normal fibroblasts.

## Results

### HCMV infection does not induce nuclear actin filaments when LifeAct-NLS is expressed at medium levels

To test if nuclear HCMV capsids would use LifeAct-stainable nuclear filamentous actin for transport, we first generated stable cell lines expressing LifeAct-mCherry-NLS similarly to an approach reported earlier (10) based on both primary HFFs (data not shown) as well as hTERT immortalized BJ cells. To visualize viral capsids, we used our recently described UL77-tagged HCMV mutant (12). Since this UL77-mGFP fusion already occupied the GFP channel, we exchanged GFP in the original LifeAct construct (RRID: Addgene_58467) with the red-fluorescent mCherry, resulting in cells showing a homogenous nuclear LifeAct signal which was slightly enriched in what seemed to be nucleoli. To our surprise, infection with either WT, UL77-GFP, or another fluorescent virus, that produces UL32EGFP and UL100mCherry, did not result in the formation of nuclear LifeAct-positive filaments in the vast majority of cells. Only very rarely (2.33% of infected cells) and only in cells expressing high amounts of LifeAct, we found nuclear filaments (Fig. 1A/B).

**Figure 1.**
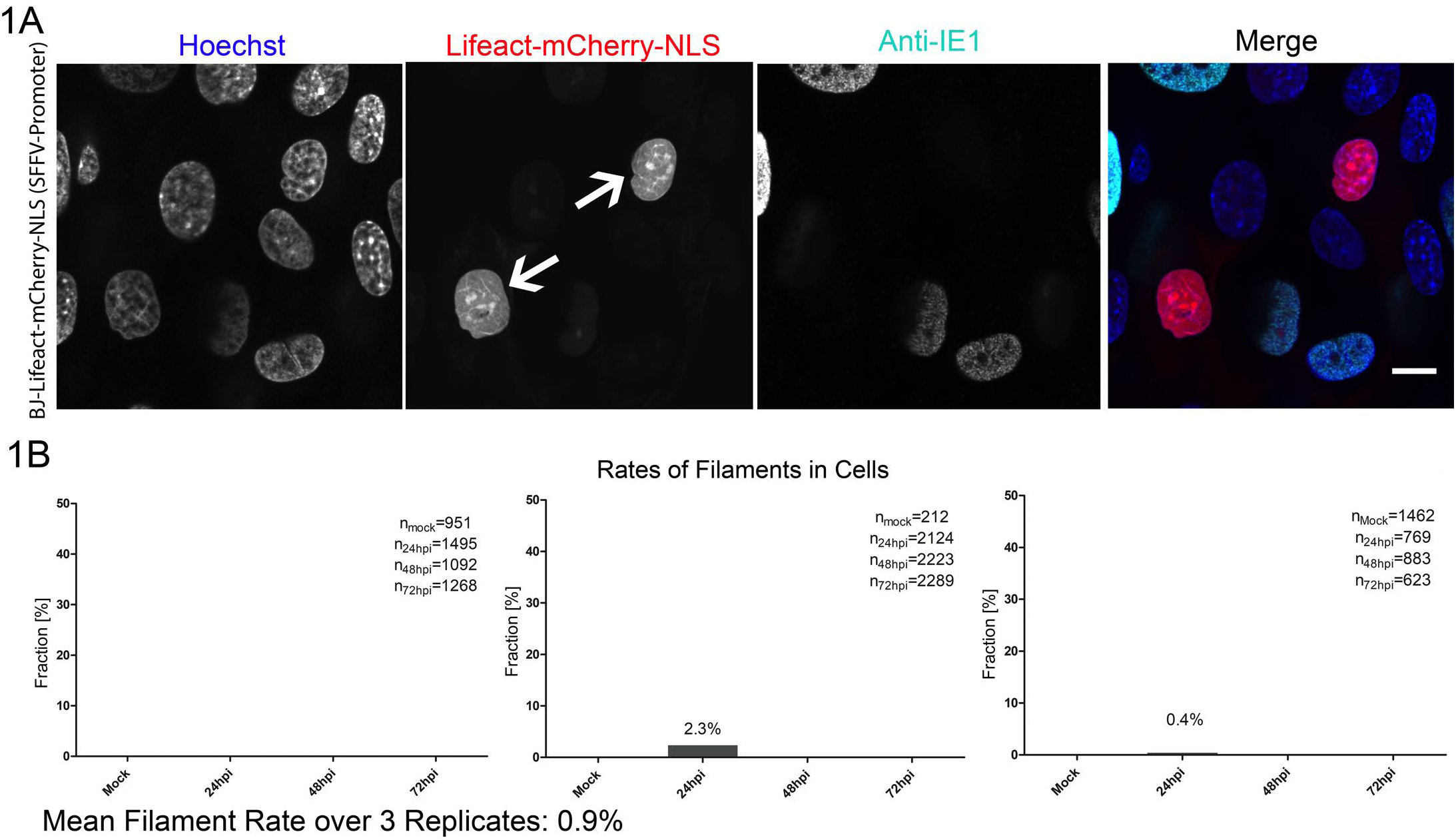
A very small fraction of cells shows nuclear F-Actin after HCMV infection. BJ-SFFV-LifeAct-mCherry-NLS cells were infected with HCMV-TB40/e-UL32EGFP-UL100mCherry, fixed at 24hpi and stained for DNA (Hoechst) and HCMV-IE (Anti-IE1). **(A)** Representative image of very rare nuclear actin filaments. Scale bar indicates 10 µm. **(B)** Quantification of the filament rates. Large tiles spanning 0.75×0.75 µm were acquired and quantified in ImageJ and Python using scripts (see Materials and Methods). Filaments were manually counted. Means of three independent replicates are shown. Bars indicate standard deviations.

### The occurrence of nuclear actin filaments correlates with the expression level of Life-Act-NLS

Interestingly, the already low frequency of nuclear filaments decreased even further with on-going infection, such that it was not possible to detect any cells with nuclear filaments later than 24 hours post infection (HPI) (Fig. 1B). We quantified nuclear LifeAct-mCherry-NLS intensities and found that intensities decreased with ongoing infection (as indicated by nuclear IE1 and nuclear pUL32-EGFP signals; Fig. 2A-C) to levels that were still easily detectable using standard excitation levels but seemingly insufficient for filament formation.

**Figure 2:**
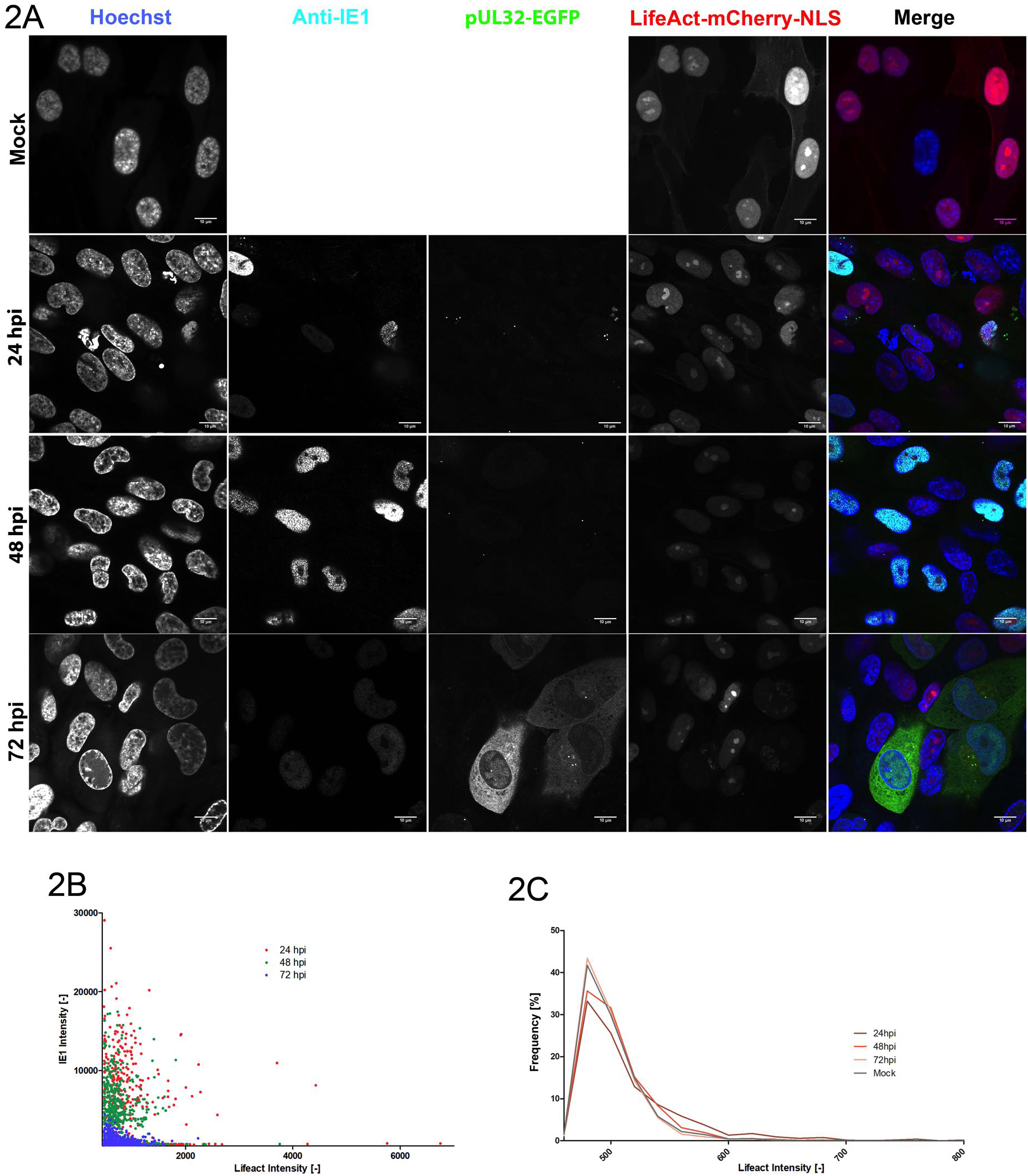
LifeAct-mCherry-NLS signal diminishes with ongoing infection. BJ-SFFV-Life-Act-mCherry-NLS cells were infected with HCMV-TB40/e-UL32EGFP-UL100mCherry, fixed at the indicated time points and stained for DNA (Hoechst) and HCMV-IE (Anti-IE1). pUL32-EGFP serves as a marker for late gene expression. **(A)** LifeAct-mCherry-NLS signal intensity drops with ongoing infection. Scale bars indicate 10 µm. **(B)** Quantification of subcellular Life-Act vs. IE signal intensities at 24, 48 and 72hpi using automated microscopy image analysis. **(C)** LifeAct intensity over time. For comparison, all image intensities are scaled to the same level in Fig. 2 and 3. Scale bars indicate 10 µm.

Since the induction of nuclear actin assemblies appeared to be dependent on high expression levels of LifeAct, we assumed that the promoter driving LifeAct-mCherry-NLS expression might critically influence the appearance of these structures. In our expression system, we used a spleen focus-forming virus-(SFFV-) promoter instead of a phosphoglycerate kinase-(PGK-) promoter utilized in the original report (10), which led to the decrease in LifeAct-mCherry-NLS signal intensity. We hypothesized that HCMV infection might interfere with expression from the SFFV promotor and therefore generated an alternative stable cell line expressing LifeAct-mCherry-NLS driven by the HCMV-Immediate early (HCMV-IE) promoter. We expected this promotor to increase LifeAct-mCherry-NLS expression in HCMV-infected cells. Indeed, we found that nuclear mCherry intensities increased when these cells were infected with HCMV up to 48hpi as shown in Fig. 3 A and B.

**Figure 3.**
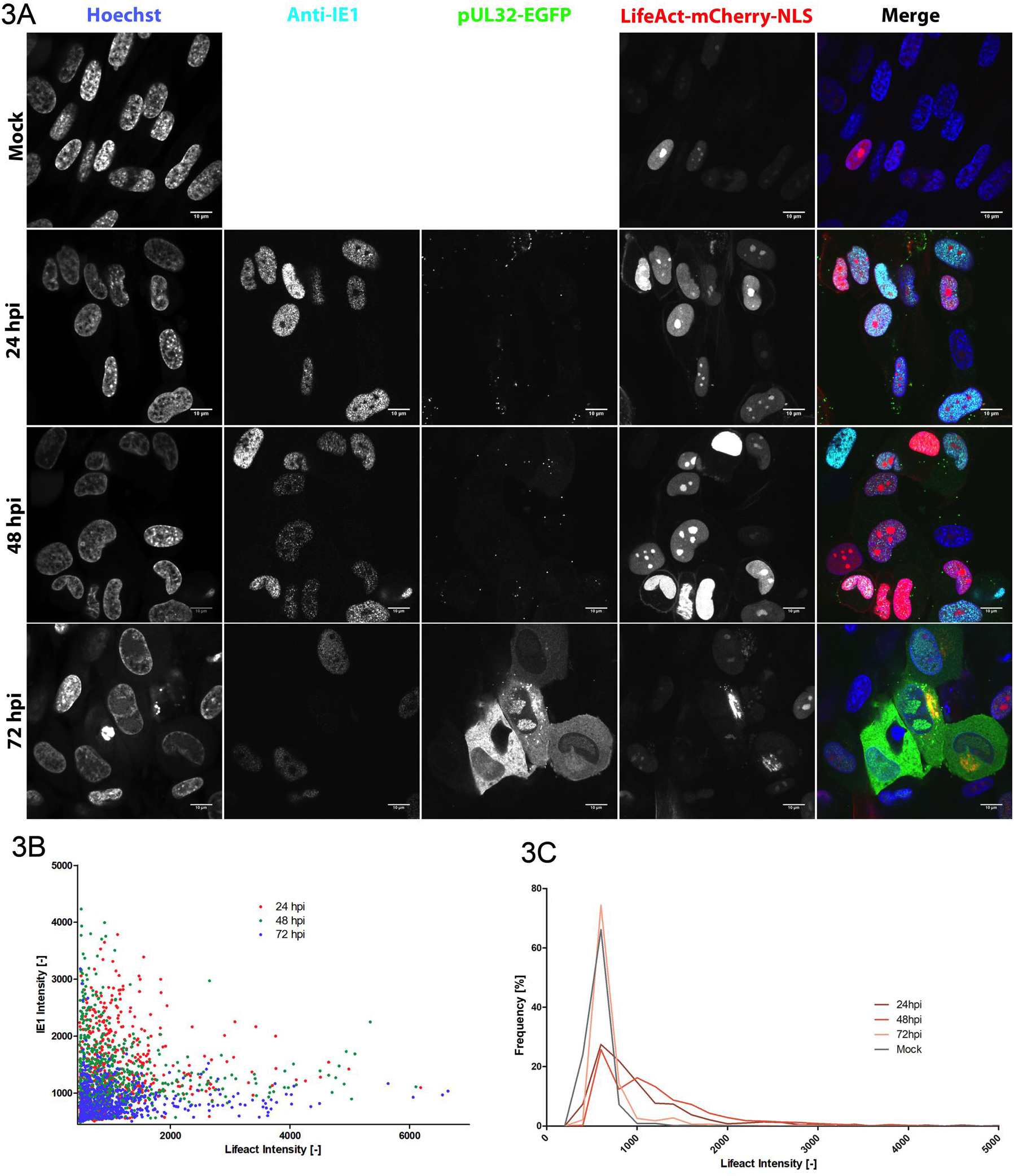
Changing the promoter of the LifeAct expression constructs alters expression dynamics in infection. BJ-CMV-LifeAct-mCherry-NLS cells were infected with HCMV-TB40/e-UL32EGFP-UL100mCherry, fixed at the indicated time points and stained for DNA (Hoechst) and HCMV-IE (Anti-IE1). pUL32-EGFP serves as a marker for late gene expression. **(A)** LifeAct-mCherry-NLS signal intensity drops with ongoing infection. Scale bars indicate 10 µm. **(B)** Quantification of subcellular LifeAct vs. IE signal intensities at 24, 48 and 72hpi by automated microscopy image analysis. **(C)** LifeAct intensity over time. All image intensities are scaled to the same level in Fig. 2 and 3 for comparison. Scale bars indicate 10 µm.

Importantly, we were now also able to detect nuclear filaments in infected cells although at lower levels compared to the earlier report (10). However, LifeAct expression decreased with progressing infection and at 72hpi reached levels that were comparable to mock-infected cells (see histograms). This effect correlated with HCMV immediate early protein 1 (IE1) expression kinetics as also shown Figs 3A and B. We did not detect filaments in the mock-infected cells, which is consistent with our previous experiments that showed no actin filaments stainable with LifeAct or with phalloidin in non-infected cells. Upon infection, we were able to visualize nuclear actin assemblies in significant quantities (Figure 4). The high expression levels of LifeAct-mCherry-NLS seemed to affect cell growth, as overall LifeAct-mCherry-NLS expression levels were reduced quickly after a few passages, which made it challenging to keep expression levels constant in between experiments. Therefore, we show the results of experimental replicates separately in Figure 4 A-D. As can be seen in Figure 4E, later passages used in experiment 2 and 3 showed reduced LifeAct-mCherry-NLS expression compared to earlier passages (experiment 1 and 4). LifeAct-mCherry-NLS expression in mock-infected cells correlated well with the amount of nuclear filament induction after infection (compare Fig. 4A/D to 4B/C), which indicates that filament induction is dependent on the LifeAct-mCherry-NLS level. Also, LifeAct-mCherry-NLS intensities were exceptionally strong in cells that showed filaments after infection compared to cells that did not as shown in Fig.4F, supporting the idea that LifeAct-mCherry-NLS acts as a concentration-dependent inducer of nuclear F-actin assembly after HCMV infection.

**Figure 4.**
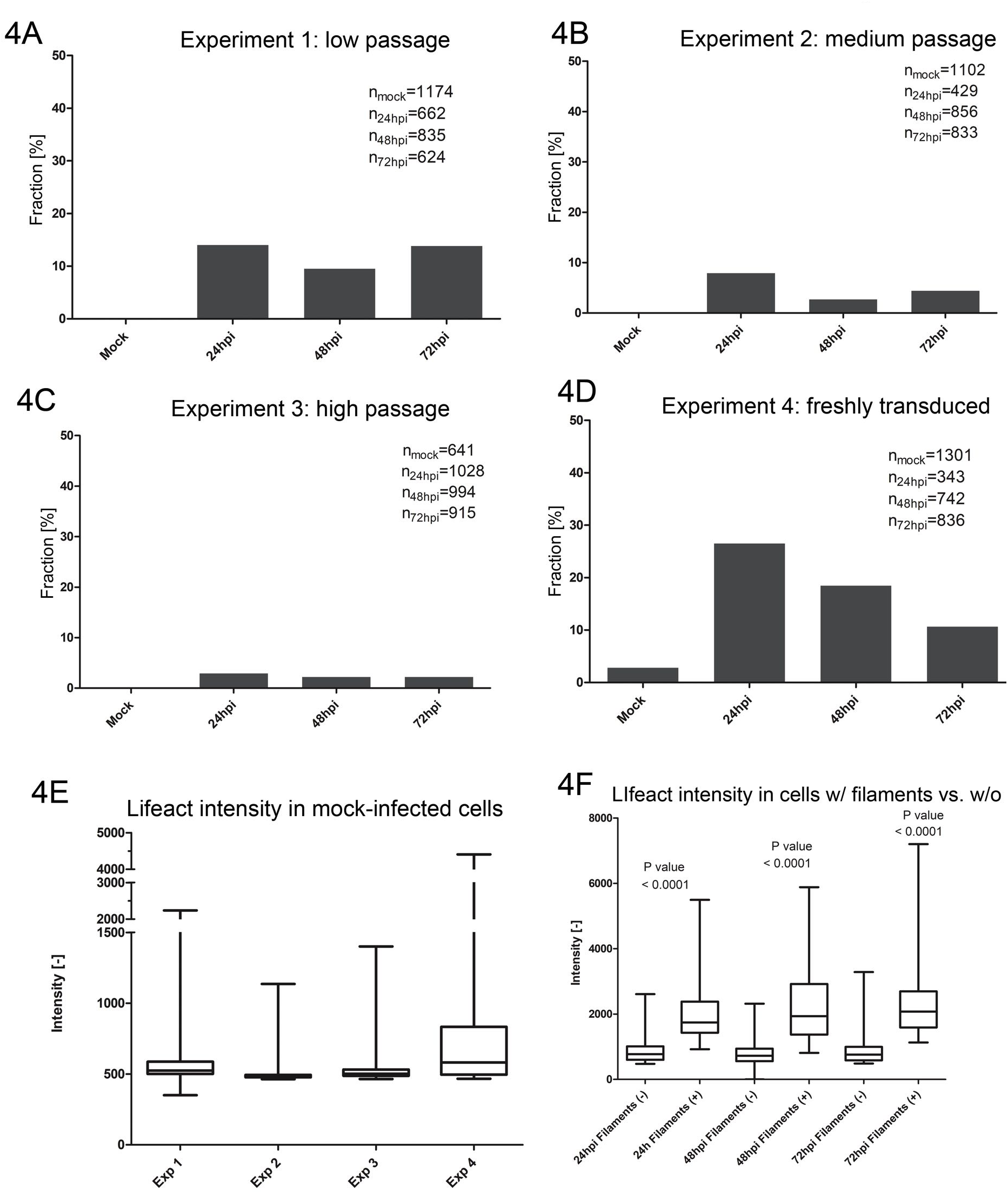
Induction of nuclear filamentous structures is dependent on LifeAct expression. BJ-CMV-LifeAct-mCherry-NLS cells were infected with HCMV-TB40/e-UL32EGFP-UL100mCherry, fixed at the indicated time points and stained for DNA (Hoechst) and HCMV-IE (Anti-IE1). **(A-D)** Rate of LifeAct nuclear filaments in infected IE-1-positive cells. Four replicates with different base-line expression levels of LifeAct are shown as quantified in **(E). (F)** Difference in LifeAct-mCherry-NLS signal intensity in the cells with filaments (+), compared to those without (-).

Of note, cells that showed nuclear actin filaments 72hpi seemed to be delayed in the progression of infection as indicated by the expression level of pUL32-EGFP (Fig. 5A, arrows), often leading to the mutual exclusion of high LifeAct and pUL32 expression (Fig. 5B).

**Figure 5.**
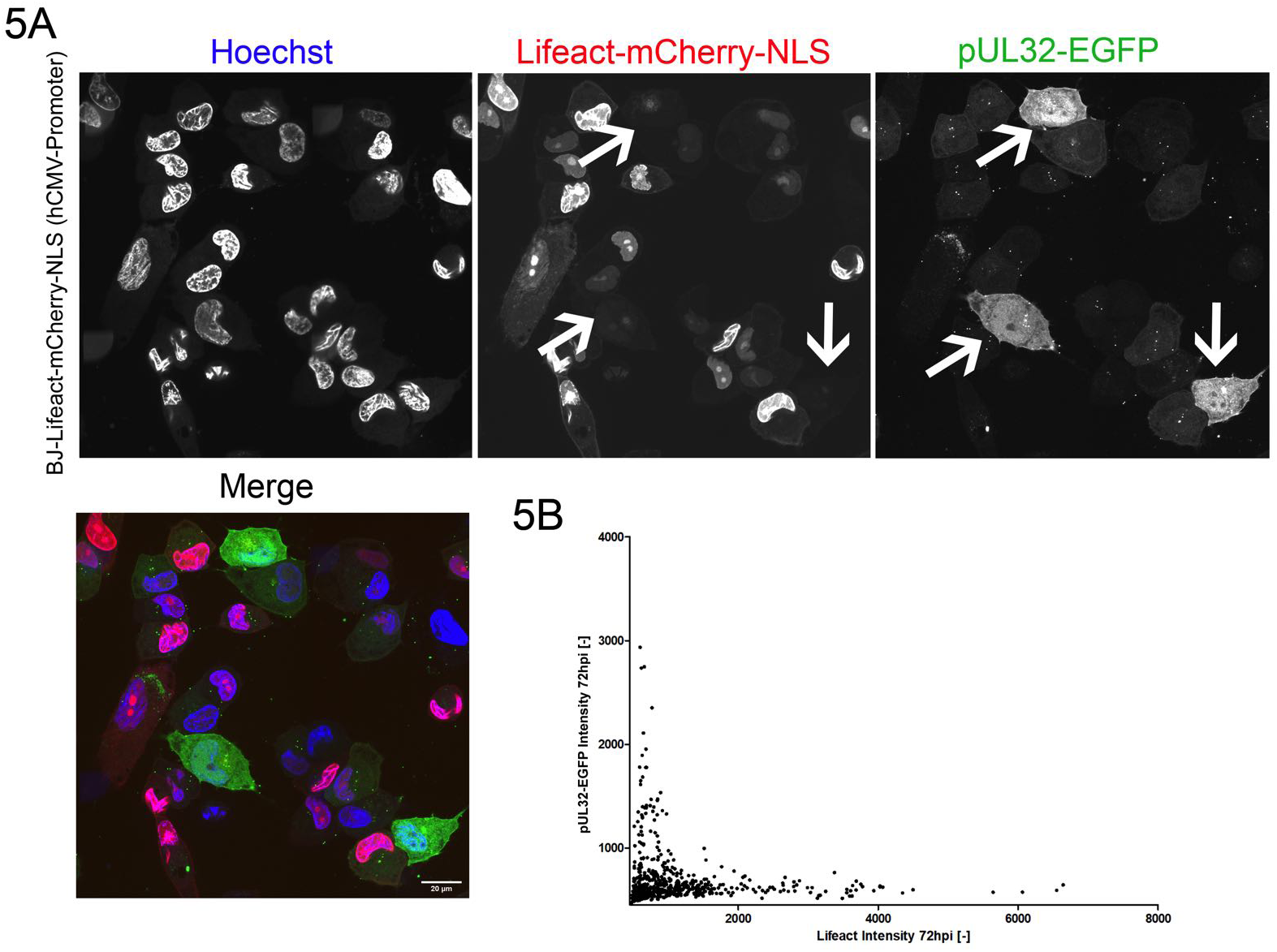
Strong LifeAct-mCherry-NLS expression and filaments induction block progress of infection. BJ-CMV-LifeAct-mCherry-NLS cells were infected with HCMV-TB40/e-UL32EGFP-UL100mCherry, fixed at 72hpi and stained for DNA (Hoechst) and HCMV-IE (Anti-IE1). pUL32-EGFP serves as a marker for late gene expression. **(A**) Cells that show pUL32-EGFP expression have very little LifeAct-mCherry-NLS signal. Scale bar indicates 20 µm. **(B)** Scatter plot of nuclear LifeAct-mCherry-NLS vs. pUL32-EGFP signal compared to the nuclear mCherry signal.

### Cells not expressing LifeAct do not show Phalloidin-stainable nuclear actin assemblies during infection

Since our results suggested that LifeAct is an inducer and also detector of filamentous nuclear actin at high expression levels, we next wanted to test if mock cells also show F-actin induction after HCMV infection. To do so, we first tested if the widely utilized fluorescent Phalloidin can detect LifeAct-induced nuclear filaments. We infected BJ cells expressing LifeAct-mCherry-NLS driven by the HCMV major immediate early promotor (MIEP) with HCMV-HB5-UL77-mGFP and fixed and stained them with fluorescent Phalloidin at 24hpi. As shown in Fig. 6, LifeAct-positive nuclear filaments could also be detected with fluorescent Phalloidin. Importantly, however, we could not detect any nuclear actin structures in wt BJ cells infected with HCMV-HB5-UL77-mGFP.

**Figure 6.**
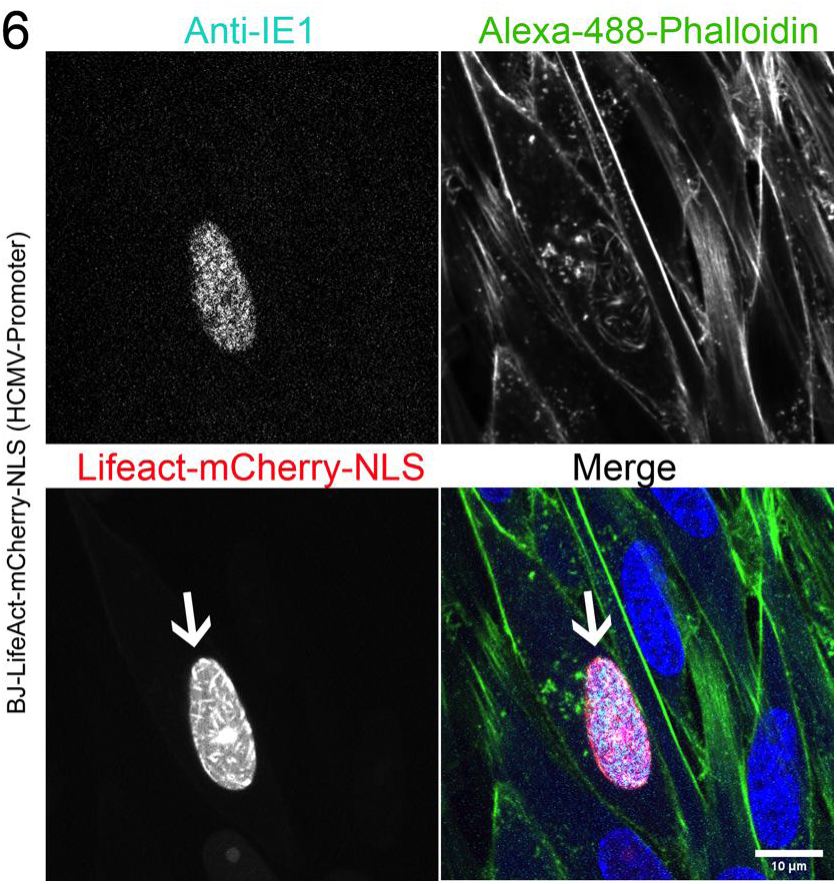
LifeAct-stained filamentous structures are detectable with Phalloidin. WT BJ and BJ-CMV-LifeAct-mCherry-NLS were infected with HCMV-HB5-UL77-mGFP at an MOI of 10 for 24 hours, fixed, and stained for IE1, as well as with Alexa-488-Phalloidin. The arrow indicates a representative BJ-CMV-LifeAct-mCherry-NLS cell in which the same nuclear actin structures are stained by both LifeAct as well as by Phalloidin. Scale bar is 10 µm.

Interestingly, almost all (95%) BJ-CMV-LifeAct-mCherry-NLS cells that showed nuclear filaments were infected as shown by IE1 staining, which indicates that the infection may be the cause of the filament induction. Again we observed that cells showing nuclear actin assemblies had exceptionally high LifeAct expression levels. These findings point towards a concentration-dependent interference of LifeAct-NLS probes with nuclear actin polymerization dynamics which has been described previously (13, 14). We also observed that the infection rate in BJ-CMV-LifeAct-mCherry-NLS cells was significantly lower than in WT-BJ cells, which further indicates that nuclear LifeAct-mCherry-NLS inhibits HCMV replication.

### Deleting the NLS precludes LifeAct-dependent induction of nuclear actin assemblies

Since our results argue for a cumulative effect of both LifeAct-mCherry-NLS expression and HCMV infection on nuclear actin availability and polymerization dynamics, we wanted to test if the NLS is the key to the induction of nuclear actin assemblies. We therefore created a cell line expressing LifeAct-mCherry without an NLS. As shown in Fig. 7, we did not detect nuclear actin filaments throughout infection in these cells, indicating that LifeAct-NLS-mediated shuttling of G-actin into the nucleus might play a role in the formation of nuclear filaments. Off note, we sometimes observed thick cytoplasmic actin assemblies in infected cells expressing a high level of LifeAct, which might indicate that LifeAct can also induce similar assemblies in the cytoplasm (Fig. 7 arrow) when convoluted with other actin-modulating interferences like HCMV infection.

### An alternative live-cell actin probe fails to detect nuclear actin assemblies during infection

Recent reports have shown that LifeAct fused to an NLS can interfere with nuclear actin dynamics (13) and some authors recommend nanobody technology as a better alternative. Especially an anti-nuclear-Actin Chromobody was referred to as minimally interfering with actin dynamics and therefore more suited to assess actin filament formation in the nucleus (13, 15). We created a mixed cell clone expressing this nuclear-Actin Chromobody to have a spectrum of different expression levels to investigate nuclear actin formation upon infection with HCMV. As shown in Figure 8, we did not find any nuclear filaments or other actin assemblies in these cells.

**Figure 7.**
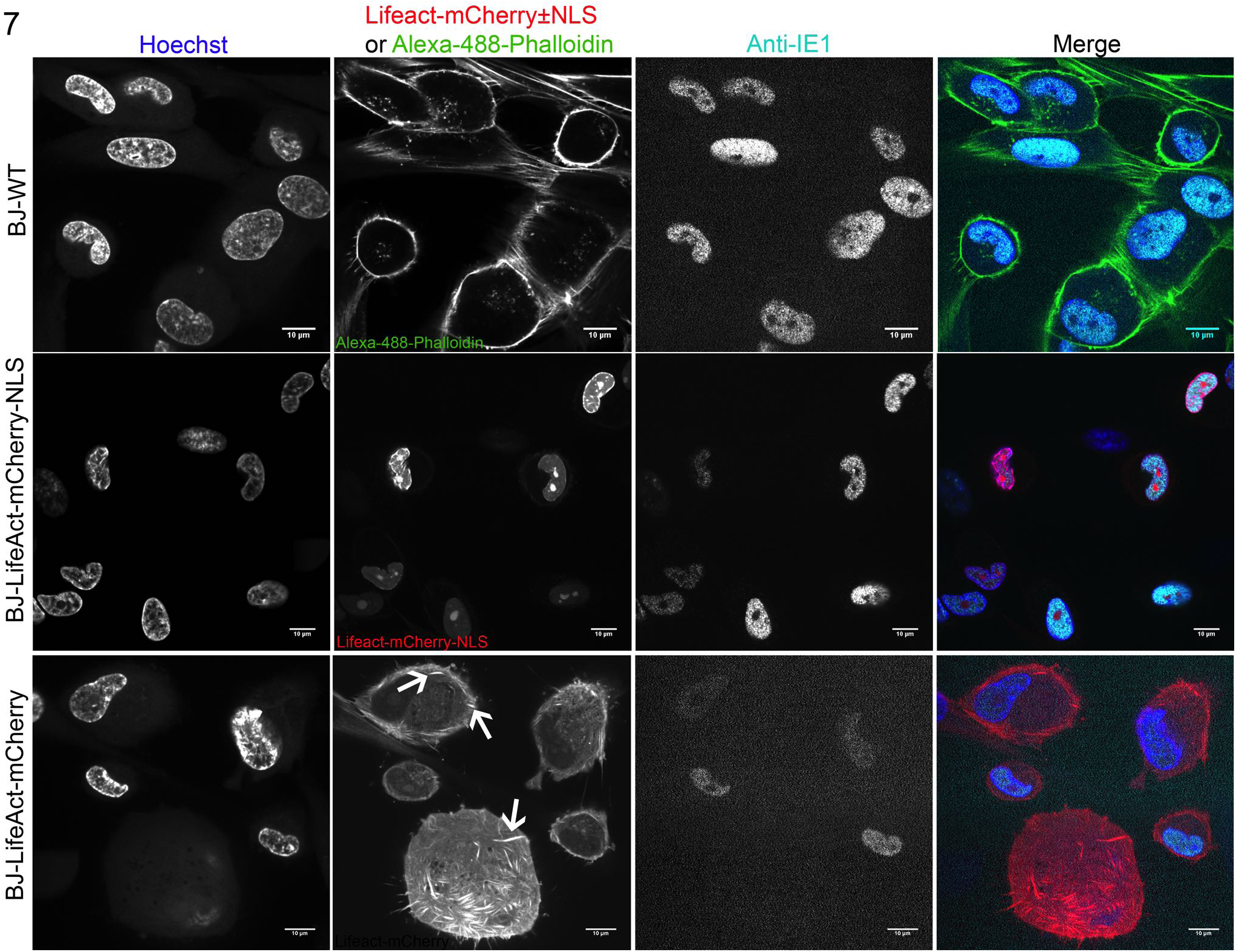
A LifeAct-mCherry fusion missing the NLS does not induce nuclear filaments. BJ-CMV-LifeAct-mCherry, WT BJ and BJ-CMV-LifeAct-mCherry-NLS were infected with HCMV-HB5-UL77-mGFP at an MOI of 10 for 24 hours, fixed, and stained for IE1, as well as with Alexa-488-Phalloidin. **(A)** No nuclear LifeAct-mCherry-stainable structures were observed without the NLS. For comparison, infected WT-BJ cells stained with Alexa-488-Phalloidin, and BJ-CMV-LifeAct-mCherry-NLS are shown. Scale bars indicate 10 µm.

**Figure 8.**
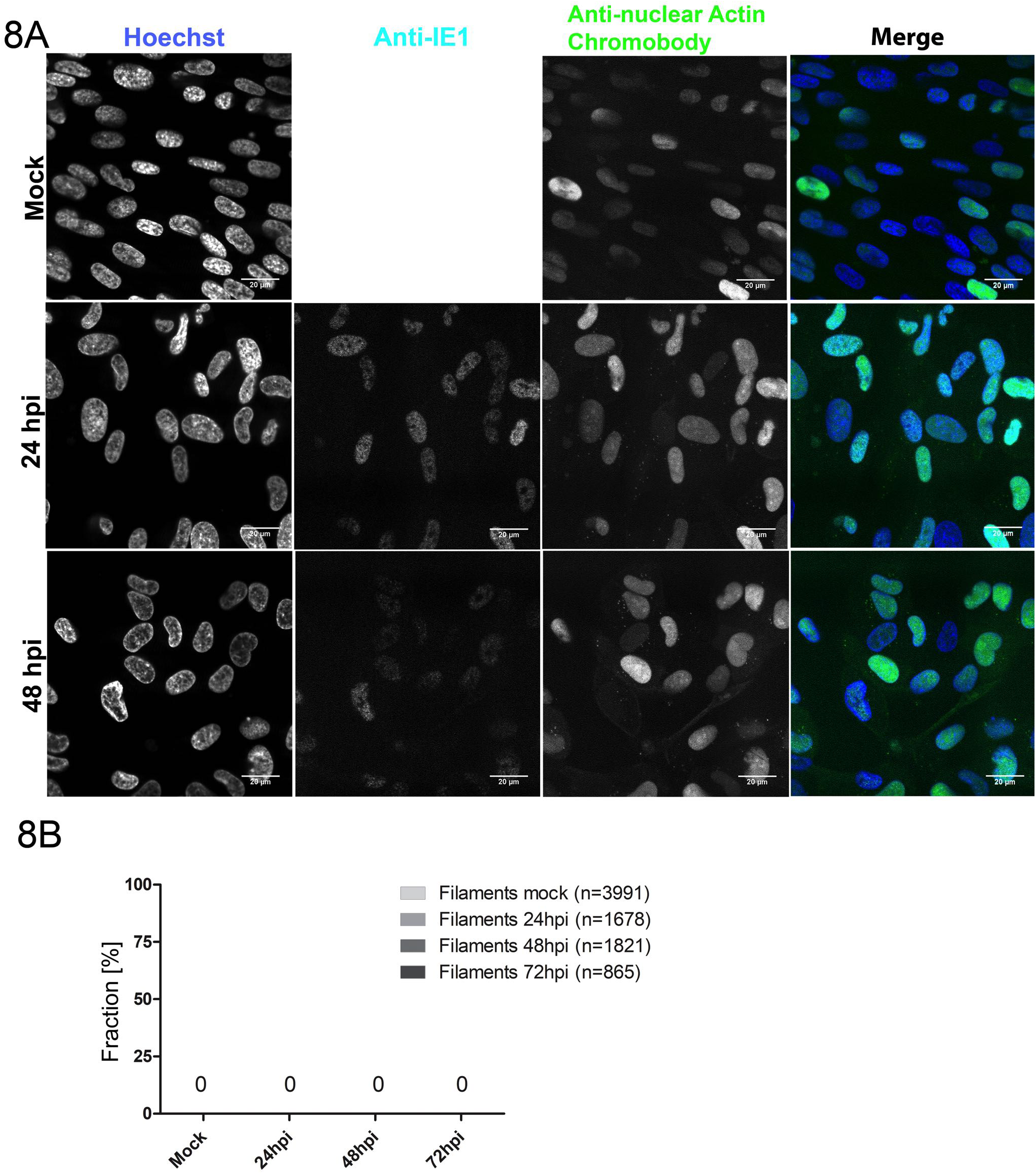
A nuclear anti-actin chromobody did not to detect nuclear actin structures. **(A)** BJ cells stably expressing a nuclear anti-actin chromobody were infected with HCMV-HB5-UL77-mGFP at an MOI of 10 for 24, 48 and 72 hours, fixed and stained for IE1. **(A)** Representative images illustrating that the chromobody does not detect any nuclear actin filaments in infected cells. Scale bars indicate 20 µm. **(B)** Quantification of filament occurrence as detected by the chromobody. One representative experiment out of 3 replicates is shown.

### Nuclear capsids do not move along LifeAct positive nuclear actin filaments

To functionally test if HCMV-induced nuclear actin filaments or actin assemblies are used for directed nuclear capsid transport, we used an HCMV mutant that produces fluorescent capsids (HCMV UL77-GFP (12)). We infected BJ cells stably expressing LifeAct-mCherry-NLS with HCMV UL77-GFP and imaged cells 72 hpi post infection. As described above, infected cells showing nuclear actin structures were extremely rare. As shown in Figure 9 and suppl. videos S1 and S2, capsids visually did not move along nuclear actin structures but instead moved in a random-walk-like fashion through the nucleus.

**Figure 9.**
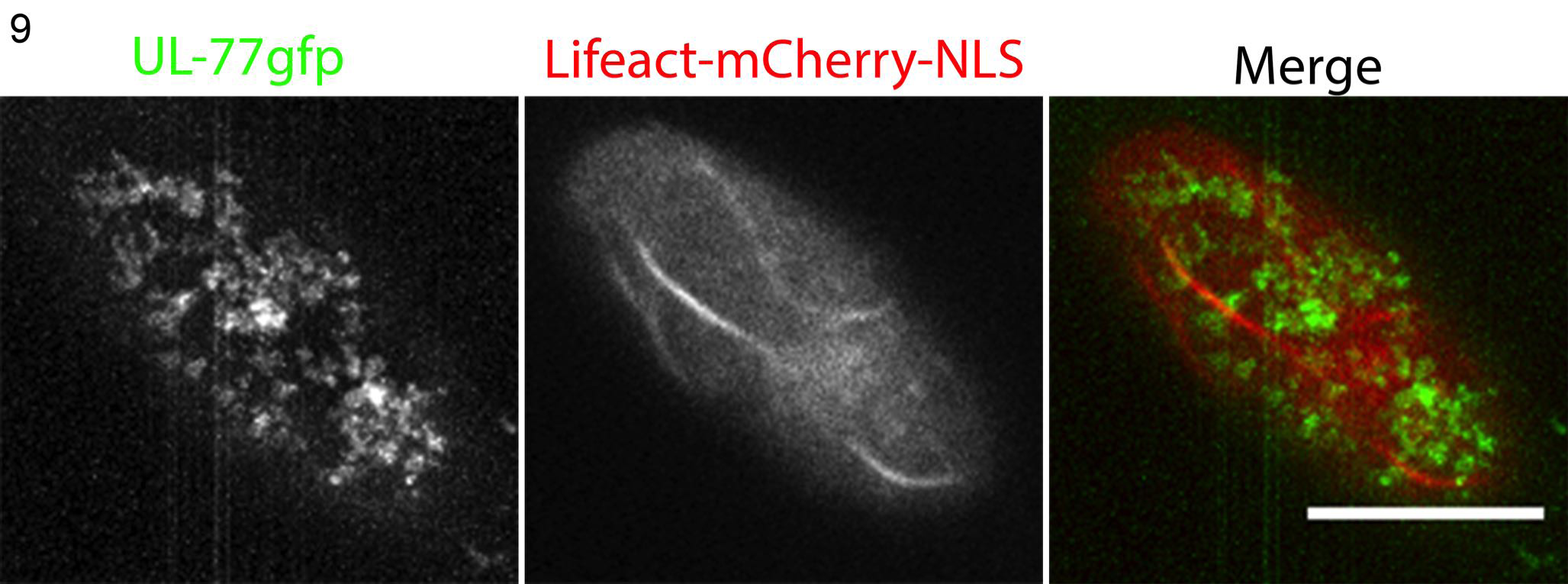
Capsids move independently of nuclear actin structures. BJ-CMV-LifeAct-mCherry-NLS cells were infected with HCMV-HB5-UL77-mGFP and imaged live at 72hpi at a frame rate of 21.45 fps. A maximum temporal projection of the GFP channel over 600 frames shows diffusive green clouds of particle location. The video is available as supplementary video S1. Scale bar indicates 10 µm.

To determine the motility mode of HCMV nuclear capsids at a quantitative level, we used single particle tracking as done previously for alphaherpesviruses (6). To this end, we infected BJ cells with HCMV UL77-GFP and imaged the cells shortly after first capsids appeared at 72 hpi. To determine particle motility modes, we optimized our established particle tracking workflow (7) using a batch-adopted version of Trackmate (16) (see Materials and Methods). Custom Matlab scripts (see Materials and Methods) allowed us to convert the data and feed it into MSDanalyzer (17) as described earlier (7).

We were able to extract a significantly larger number of particle tracks from our data using this approach, increasing the statistical validity of our analysis. In line with our visual assessment, our quantification of more than 17000 single tracks longer than 1 second (20 frames) showed that HCMV nuclear capsids do not engage in directed motility, but show slight sub-diffusion with an anomalous diffusion exponent α of 0.74 in the nucleoplasm over short time-scales (Fig. 10). This result is a little lower than our previously published data for HSV-1 and PRV. A calculation of corral size, performed as in (7), revealed a corral size of approximately 700nm and indicates that also HCMV remodels the nuclear structure very similarly to HSV-1 and PRV, which in turn would allow nuclear capsids to cross large areas of the nuclear space by diffusion.

**Figure 10.**
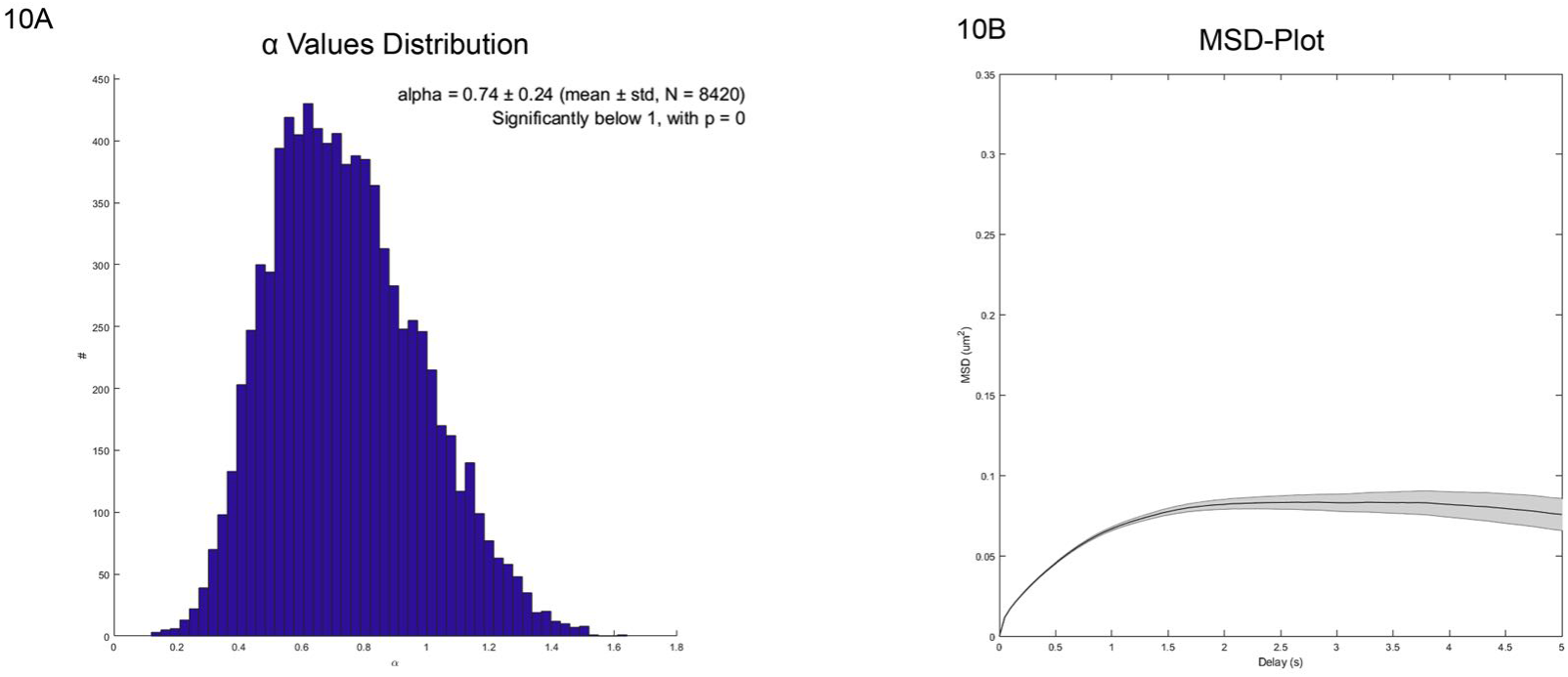
Single particle tracking reveals diffusion as the major nuclear capsid motility mode. WT BJ cells were infected with HCMV-HB5-UL77-mGFP and imaged live at 72hpi at a frame rate of 21.45 fps, and capsids were tracked using a custom batch version of Trackmate. Tracks were subsequently analyzed utilizing MSD Analyzer, and the diffusion exponent alpha **(A)**, as well as the average corral size using an MSD plot **(B)**, were extracted. The saturation of the MSD curve at about 0.08 indicates a chromatin corral diameter of about 700 nm.

### Electron microscopy could not to detect nuclear actin bundles in normal infected cells

To determine if normal infected cells that do not express LifeAct-NLS show any form of nuclear filaments that cannot be detected using fluorescent probes, we applied electron microscopy as an additional tool. We seeded BJ-CMV-LifeAct-mCherry-NLS cells on sapphire discs and infected them with HCMV. The infected cells were checked by live-cell fluorescence microscopy for the appearance of nuclear actin structures and subsequently high-pressure-frozen and processed for transmission electron microscopy. Indeed, we found thick bundles of nuclear filaments in a fraction of cells expressing LifeAct-mCherry-NLS (Fig. 11A-B, S3). However, in line with our previous experiments, we could not detect nuclear filaments in cells that expressed LifeAct-mCherry-NLS at low levels. The bundles showed strong similarity to previously published nuclear actin bundles induced by overexpression of actin mutants (18). The bundles appeared more often close to the nuclear envelope, but we did not find evidence for filaments connecting the nuclear envelope to a nascent central viral replication compartment as proposed earlier (10).

**Figure 11.**
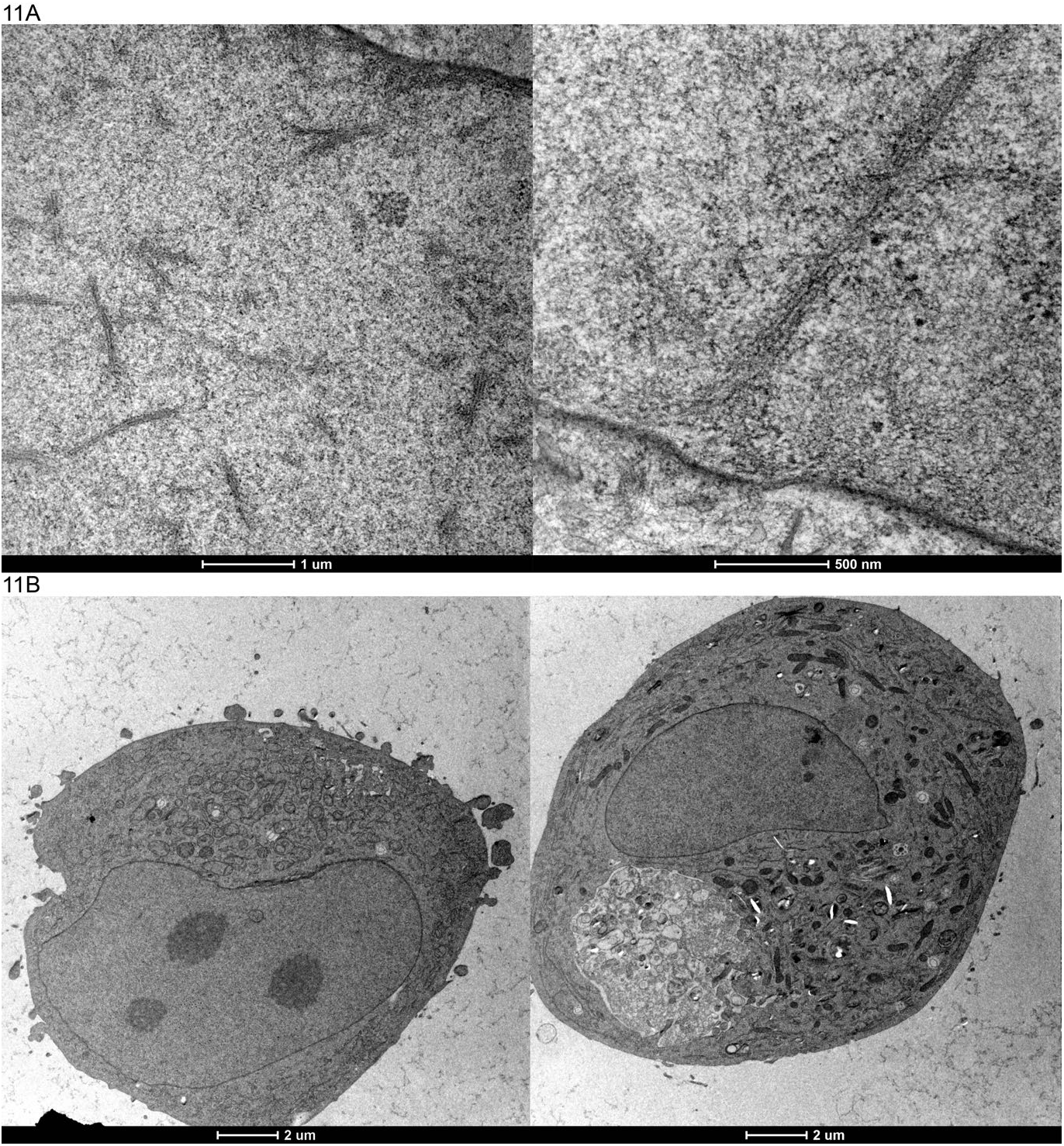
Electron microscopy only reveals thick filamentous bundles in infected cells expressing LifeAct-mCherry-NLS. BJ or BJ-CMV-LifeAct-mCherry-NLS cells were infected with HCMV-TB40/e-UL32EGFP-UL100mCherry, high-pressure-frozen at 24hpi and freeze-substituted. **(A)** Bundled, filamentous actin structures can be visualized in the nuclei of infected cells expressing LifeAct-mCherry-NLS. Bundles only appeared in a fraction of the examined cells and in higher frequency closer to the nuclear envelope (see also supplementary Figure S3). **(B)** Two cells without apparent nuclear actin structures visible for comparison. For scale bar length see picture.

## Discussion

In this study, we investigated nuclear actin filament formation after HCMV infection. We found that only infection of cells highly overexpressing LifeAct-mCherry-NLS leads to nuclear filament formation. Cells expressing lower levels of nuclear LifeAct-mCherry-NLS (which was still easily detectable) did not show filament induction. Furthermore, Phalloidin failed to detect nuclear filaments in infected cells that do not express LifeAct-mCherry-NLS, but was able to mark filaments in infected, LifeAct-overexpressing cells. We, therefore, conclude that LifeAct-mCherry-NLS is a concentration-dependent inducer of nuclear actin structures in HCMV-infected cells. Our results are supported by a previous study that showed that nuclear actin filaments could be induced by overexpression of NLS-actin fusion proteins (18) as well as a more recent study that describes the interference of LifeAct-NLS probes with nuclear actin dynamics, resulting in the formation of nuclear actin assemblies (13).

For this reason, a recent review recommended the use of alternative markers to assess the nuclear actin dynamics in living cells such as an anti-actin-chromobody (15). This probe failed to detect any filamentous actin structures in normal, HCMV-infected cells. In line with these results, TEM analysis showed large nuclear bundled filamentous structures in a subpopulation of HCMV-infected cells that expressed high levels of LifeAct-NLS. These structures are easily detectable in EM if they appear in normal infected cells. However, we could not detect any of those structures in normal infected BJ cells. To our knowledge, there is also no EM-based evidence of nuclear actin bundles in HCMV-infected cells in the literature.

In our hands, capsid formation and nuclear filament induction appeared to be almost mutually exclusive, which indicates that overexpression of the LifeAct-mCherry-NLS probe is detrimental for the progression of viral infection. For this reason, we could visualize capsid movement in the presence of actin assemblies in a few cells only. Still, capsid movement was independent of actin assemblies and was indistinguishable from capsid motility in fibroblasts not expressing LifeAct. Single particle tracking of HCMV nuclear capsids showed that diffusion is the main motility mode, and confirmed our previous results for HSV-1 and PRV (6, 7), indicating that nuclear capsid diffusion might be a conserved motility mode to cross the nucleoplasm. Support for this conclusion also comes from two recent studies in which the authors first show that HSV-1 induces egress channel formation through the marginalized chromatin (8), which could represent the motility spaces that can be measured by tracking nuclear capsids ((7) and Fig. 10). Secondly, computational simulations using our experimentally determined diffusion coefficient suggest that these egress channels allow capsid translocation to the nuclear membrane by diffusion within minutes (9).

The question remains which molecular mechanism induces filament formation in LifeAct-NLS-expressing cells. Since a LifeAct construct without the NLS sequence was unable to induce actin structures in the nucleus of infected cells, we hypothesize that LifeAct-NLS increases the concentration of monomeric G-actin in the nucleus. Infection-induced disruption of cytoplasmic F-actin (19) might increase the available G-actin pool, which could result in exceeding a nuclear concentration threshold at which filament formation occurs (18). While Wilkie et al. (10) did not detect an increase of nuclear actin monomers in infected cells, they did not compare cells with and without the LifeAct-GFP-NLS construct. It is therefore currently not clear if increased amounts of monomeric G-actin get transported into the nucleus by LifeAct-NLS during HCMV infection.

The role of actin in the nucleus is a matter of current debate, and intimately connected to a discussion about the strengths and weaknesses of the actin probes used (14, 15, 20–25). A variety of roles has been described for nuclear actin in recent years, ranging from gene regulation to structural organization (24). Widely used probes like LifeAct can interfere with nuclear actin polymerization, and these caveats must be taken into account when assessing the biological role of nuclear actin filaments (13, 20). The reasons for the artifacts are often unclear. However, it is likely that nuclear-targeted probes alter the nuclear concentration of actin monomers by shuttling G-actin into the nucleus (13, 14), which is also supported by our results. Probe-induced nuclear filament formation has not only been described for LifeAct-NLS, but also for UTR261 and UTR230-NLS fusions (13, 14).

Based on our results we conclude that HCMV does not induce large-scale nuclear actin assemblies, and that nuclear capsid motility is not dependent on large actin tracks. However, we cannot exclude that more subtle and maybe transient actin assemblies play a role in nuclear morphogenesis events. Possible targets might be genome encapsidation and release of capsids from the replication compartment. Future studies will have to be carefully designed to examine the role of actin in nuclear herpesvirus morphogenesis while omitting the known pitfalls of nuclear actin probes.

## Materials and methods

### Cells and viruses

BJ-5ta hTERT-immortalized human fibroblasts were licensed from ATCC (CRL-4001) and cultivated in Dulbecco’s Modified Eagles Medium Glutamax^®^ (Thermofisher) with 20% Medium 199 (Earles Salts) (Thermofisher), 10% FBS superior (Merck), 0.8 mM sodium pyruvate (Thermofisher) and 1 µg/ml Hygromycin (Invivogen).

BAC-derived AD169-based HCMV-HB5-UL77-mGFP is described in reference (12). HCMV-TB40-BAC_kl.7_-UL32EGFP-UL100mCherry is described in Sampaio et al. (26). The also TB40 based HCMV-UL32-EGFP is reported in (27).

### pSFFV and pCMV-driven LifeAct-2XNLS expression constructs

The lentiviral plasmid LeGO_SFFV_LifeAct-mCh-2XNLS was generated by designing a LifeAct-mCherry-2XNLS insert reflecting the pEGFP-C1_LifeAct-EGFP_2XNLS construct (20). (pEGFP-C1 LifeAct-EGFP-2XNLS was a gift from Dyche Mullins (RRID: Addgene_58467). We also introduced an upstream 5’ AvrII site to facilitate cloning. The resulting sequence was inserted into LeGO-iC2 (28), thereby replacing the original IRES-mCherry sequence of LeGO-iC2 between BamHI and BsrGI. To generate LeGO_CMV_LifeAct-mCh-2XNLS, we replaced the SFFV promoter of the original gene expression cassette by the HCMV major immediate early pro-moter sequence via standard PCR-based cloning using NheI and AvrII. Correct sequences of both expression constructs were confirmed by Sanger sequencing.

### Immunofluorescence

For quantification of LifeAct or chromobody intensities during HCMV infection, cells were cultivated in BJ-Medium and seeded on Ibidi glass bottom µ-Dishes coated with Fibronectin 1:100 in Dulbecco’s phosphate buffered saline (D-PBS, Sigma-Al-drich). After infection with HCMV-TB40-BAC_kl.7_-UL32EGFP-UL100mCherry or HCMV-UL32GFP (gift from Lüder Wiebusch) at a multiplicity of infection (MOI) of 10. Cells were fixed with 4% paraformaldehyde (Science Services) in D-PBS at 24hpi, 48hpi, and 72hpi. For immunofluorescence (IF) staining, the cells were permeabilized with 0.1% TritonX100 in D-PBS, blocked with 3% Bovine Serum Albumin (BSA) in D-PBS, and subsequently stained with a primary murine anti-IE1 antibody and a secondary Alexa 647 Goat anti-mouse antibody (Thermofisher). Nuclei were additionally stained with Hoechst 33342 (Thermofisher). For Phalloidin staining, BJ-WT and BJ-CMV-LifeAct-mCherry-NLS were infected with HCMV-HB5-UL77-mGFP at an MOI of 1 for 24h and subsequently fixed and stained with Alexa-488-Phalloidin (Thermofisher) and Hoechst 33342 (Thermofisher).

**Microscopy** was performed with a Nikon spinning disc system consisting of a Yokogawa W2 and two Andor iXON888 cameras using NIS-Elements for image acquisition. A Nikon 100x 1.49 NA Apo-TIRF objective was used resulting in 130nm pixel size. The system was equipped with standard 405, 488, 561, 640 nm laser lines and corresponding filter sets. For quantification, 5×5 image tiles were acquired, resulting in a 666x666 µm captured area.

**Image analysis** was performed with an ImageJ macro and a python Jupyter notebook (both available on github through https://github.com/QuantitativeVirology/FIJI-Segmentor-Macro and https://github.com/QuantitativeVirology/FIJI-Measurement-Analyzer). Nuclei segmentation was done in ImageJ using the Hoechst channel creating regions of interest (ROIs), which were subsequently used to measure the signal intensities in the other channels of interest. Resulting mean signal intensities were processed in Python. The number of cells containing filaments was counted using the manual cell counter plugin from ImageJ. Statistical analysis was done with GraphPad PRISM.

### Single particle tracking

For tracking of single viral particles in cell nuclei, BJ-WT cells were cultivated in BJ-Medium and infected with HCMV-HB5-UL77-mGFP (MOI of 1.5). 72hpi vid-eos of living cells were acquired with 21.45 frames per second (fps) at 37°C with 5% CO_2_. Single nuclei were cropped, and capsids were tracked with the Fiji plugin Trackmate by Tinevez et al. (16), using a custom batch analysis plugin (available on github through https://github.com/QuantitativeVirology/Trackmate_Batch). The resulting XML files were analyzed using custom Matlab scripts (available on github through https://github.com/QuantitativeVirology/Matlab-Trackmate-MSD), which in turn make use of the Matlab class “Mean square displacement analysis of particle trajectories”, also from Tinevez and colleagues (17). Visualization of the results was also done with Matlab.

### Electron Microscopy

For structural analysis of nuclear actin assemblies, Sapphire discs (M. Wohlwend) with a diameter of 3mm and a thickness of 0.17mm were pre-cleaned by immersion in soapy water and sonication for 10 minutes. Afterward, the discs were washed in >99% Ethanol (Roth) twice by sonication for 10 minutes each and plasma cleaned in a Quorum Q150 plus (Quorum Technologies Ltd, UK) machine for 120 seconds and subsequently coated with a thin film of carbon through carbon cord evaporation. The discs were dried overnight at 60°C and kept at that temperature until shortly before use.

BJ-CMV-LifeAct-mCherry-NLS cells were cultivated in BJ-Medium and seeded on the previously prepared sapphire discs. On the following day, the cells were infected with HCMV-TB40e-UL32EGFP-UL100mCherry at an MOI of 10. After 24 hours cells were imaged through live-cell spinning disc microscopy to check for LifeAct-induced filaments. Afterwards, cells were high-pressure frozen as described in (29).

For freeze-substitution, sapphire discs were incubated in -90°C pre-cooled freeze substitution medium consisting of 0.2% Osmium tetroxide (Science Services), 0.1% Uranyl acetate (Merck) and 5% water in Acetone (Merck) overnight at -90°C in an Arctiko DP-80 cryo porter (Arctiko, Denmark) and subsequently thawed by stopping the cooling and leaving the machine to warm to room temperature.

The freeze-substituted samples were subsequently embedded in Epon and cut into ultrathin 50nm slices using a Leica Ultracut microtome (Leica, Germany). The slices were transferred to copper mesh grids, post-stained in saturated Uranyl acetate in 70% Ethanol (Roth) for 7 minutes and subsequently imaged using an FEI Tecnai F20 electron microscope (Thermofisher, USA).

## Acknowledgments

We thank Christian Sinzger (University of Ulm, Germany) for the gift of dual-color HCMV through Jens von Einem (University of Ulm, Germany). We also thank Lüder Wiebusch (Charite Berlin) for HCMV-UL32GFP. pEGFP-C1 LifeAct-EGFP-2XNLS was a gift from Dyche Mullins through Addgene (plasmid # 58467). LeGO-iC2 was a kind gift of Kristoffer Riecken and Boris Fehse (UKE Hamburg). This study was funded through a Wellcome Trust collaborative award to JBB (209250/Z/17/Z) and KG (209250/Z/17/Z). The Heinrich Pette Institute, Leibniz Institute for Experimental Virology is supported by the *Free and Hanseatic City of Hamburg* and the *Federal Ministry of Health*. MM and KG are supported by the Cluster of Excellence RESIST (EXC 2155), Hannover Medical School, Hannover, Germany.

## Supplementary Material

**Video S1 and S2. Single viral particles are moving through the nucleus of an infected cell.** In these videos, the nuclei of BJ-CMV-LifeAct-mCherry-NLS cell infected with HCMV-HB5-UL77-mGFP (MOI of 1.5) are shown at 72hpi. They represent rare examples in which nuclear actin assemblies and viral capsids are visible in the nucleus. Viral particles do not show obvious movement along the filaments. Instead, they move in a random-walk like behavior through the nucleoplasm.

**Figure S3. Electron micrograph of a nucleus from an infected, high-pressure frozen cell.** Lower section of an BJ-CMV-LifeAct-mCherry-NLS cell infected with HCMV-TB40/e-UL32EGFP-UL100mCherry which was high-pressure-frozen at 24hpi and subsequently freeze-substituted. This cell is an example of the high density of actin bundles adjacent to the nuclear envelope. (A/B) Details of the bundles shown. Scales are indicated in the pictures.

## References

1. Sanchez V, Greis KD, Sztul E, Britt WJ. 2000. Accumulation of Virion Tegument and Envelope Proteins in a Stable Cytoplasmic Compartment during Human Cytomegalovirus Replication: Characterization of a Potential Site of Virus Assembly. J Virol 74:975–986.

2. Sanchez V, Sztul E, Britt WJ. 2000. Human Cytomegalovirus pp28 (UL99) Localizes to a Cytoplasmic Compartment Which Overlaps the Endoplasmic Reticulum-Golgi-Intermediate Compartment. J Virol 74:3842–3851.

3. Mettenleiter TC, Müller F, Granzow H, Klupp BG. 2013. The way out: what we know and do not know about herpesvirus nuclear egress. Cell Microbiol 15:170–178.

4. Forest T, Barnard S, Baines JD. 2005. Active intranuclear movement of herpesvirus capsids. Nature Cell Biology 7:429–431.

5. Feierbach B, Piccinotti S, Bisher M, Denk W, Enquist LW. 2006. Alpha-Herpesvirus Infection Induces the Formation of Nuclear Actin Filaments. PLoS Pathogens 2:e85.

6. Bosse JB, Virding S, Thiberge SY, Scherer J, Wodrich H, Ruzsics Z, Koszinowski UH, Enquist LW. 2014. Nuclear Herpesvirus Capsid Motility Is Not Dependent on F-Actin. Mbio 5:e01909–14.

7. Bosse JB, Hogue IB, Feric M, Thiberge SY, Sodeik B, Brangwynne CP, Enquist LW. 2015. Remodeling nuclear architecture allows efficient transport of herpesvirus capsids by diffusion. Proceedings of the National Academy of Sciences 112:E5725–E5733.

8. Myllys M, Ruokolainen V, Aho V, Smith EA, Hakanen S, Peri P, Salvetti A, Timonen J, Hukkanen V, Larabell CA, Vihinen-Ranta M. 2016. Herpes simplex virus 1 induces egress 1. channels through marginalized host chromatin. Scientific Reports 6:28844.

9. Aho V, Myllys M, Ruokolainen V, Hakanen S, Mäntylä E, Virtanen J, Hukkanen V, Kühn T, Timonen J, Mattila K, Larabell CA, Vihinen-Ranta M. 2017. Chromatin organization regulates viral egress dynamics. Sci Rep-uk 7:3692.

10. Wilkie AR, Lawler JL, Coen DM. 2016. A Role for Nuclear F-Actin Induction in Human Cytomegalovirus Nuclear Egress. mBio 7:e01254–16.

11. Wilkie AR, Sharma M, Pesola JM, Ericsson M, Fernandez R, Coen DM. 2018. A Role for Myosin Va in Human Cytomegalovirus Nuclear Egress. J Virol 92:e01849–17.

12. Borst E, Bauerfeind R, Binz A, Stephan T, Neuber S, Wagner K, Steinbrück L, Sodeik B, Roviš T, Jonjić S, Messerle M. 2016. The Essential Human Cytomegalovirus Proteins pUL77 and pUL93 Are Structural Components Necessary for Viral Genome Encapsidation. J Virol 90:5860–5875.

13. Du J, Fan Y, Chen T, Feng X. 2015. Lifeact and Utr230 induce distinct actin assemblies in cell nuclei. Cytoskeleton 72:570–575.

14. Belin BJ, Mullins DR. 2013. What we talk about when we talk about nuclear actin. Nucl Austin Tex 4:291–7.

15. Melak M, Plessner M, Grosse R. 2017. Actin visualization at a glance. J Cell Sci 130:jcs.189068.

16. Tinevez J-Y, Perry N, Schindelin J, Hoopes GM, Reynolds GD, Laplantine E, Bednarek SY, Shorte SL, Eliceiri KW. 2017. TrackMate: An open and extensible platform for singleparticle tracking. Methods San Diego Calif 115.

17. Tarantino N, Tinevez J-Y, Crowell E, Boisson B, Henriques R, Mhlanga M, Agou F, Israël A, Laplantine E. 2014. TNF and IL-1 exhibit distinct ubiquitin requirements for inducing NEMO–IKK supramolecular structures. J Cell Biology 204:231–245.

18. Kokai E, Beck H, Weissbach J, Arnold F, Sinske D, Sebert U, Gaiselmann G, Schmidt V, Walther P, Münch J, Posern G, Knöll B. 2014. Analysis of nuclear actin by overexpression of wild-type and actin mutant proteins. Histochemistry and Cell Biology 141:123–135.

19. Jones N, Lewis J, Kilpatrick. 1986. Cytoskeletal disruption during human cytomegalovirus infection of human lung fibroblasts. Eur J Cell Biol 41:304–12.

20. Belin BJ, Cimini BA, Blackburn EH, Mullins DR. 2013. Visualization of actin filaments and monomers in somatic cell nuclei. Molecular Biology of the Cell 24:982–994.

21. Courtemanche N, Pollard TD, Chen Q. 2016. Avoiding artefacts when counting polymerized actin in live cells with LifeAct fused to fluorescent proteins. Nat Cell Biol 18:676–683.

22. Munsie LN, Caron N, Desmond CR, Truant R. 2009. Lifeact cannot visualize some forms of stress-induced twisted f-actin. Nat Methods 6:nmeth0509-317.

23. Riedl J, Crevenna AH, Kessenbrock K, Yu J, Neukirchen D, Bista M, Bradke F, Jenne D, Holak TA, Werb Z, Sixt M, Wedlich-Soldner R. 2008. Lifeact: a versatile marker to visualize F-actin. Nat Methods 5:605–607.

24. Viita T, Vartiainen MK. 2016. From Cytoskeleton to Gene Expression: Actin in the Nucleus. Handb Exp Pharmacol.

25. Flores LR, Keeling MC, Zhang X, Sliogeryte K, Gavara N. 2019. Lifeact-GFP alters Factin organization, cellular morphology and biophysical behaviour. Sci Rep-uk 9:3241.

26. Sampaio K, Jahn G, Sinzger C. 2013. Virus-Host Interactions, Methods and Protocols. Methods Mol Biology Clifton N J 1064:201–209.

27. Sampaio K, Cavignac Y, Stierhof Y-D, Sinzger C. 2005. Human Cytomegalovirus Labeled with Green Fluorescent Protein for Live Analysis of Intracellular Particle Movements. J Virol 79:2754–2767.

28. Weber K, Bartsch U, Stocking C, Fehse B. 2008. A Multicolor Panel of Novel Lentiviral “Gene Ontology” (LeGO) Vectors for Functional Gene Analysis. Mol Ther.

29. Höhn K, Sailer M, Wang L, Lorenz M, Schneider ME, Walther P. 2011. Preparation of cryofixed cells for improved 3D ultrastructure with scanning transmission electron tomography. Histochem Cell Biol 135:1–9.

