## Supplementary figures and images for "Human cytomegalovirus nuclear capsid motility is non-directed and independent of nuclear actin bundles"

### supplemental figure S3

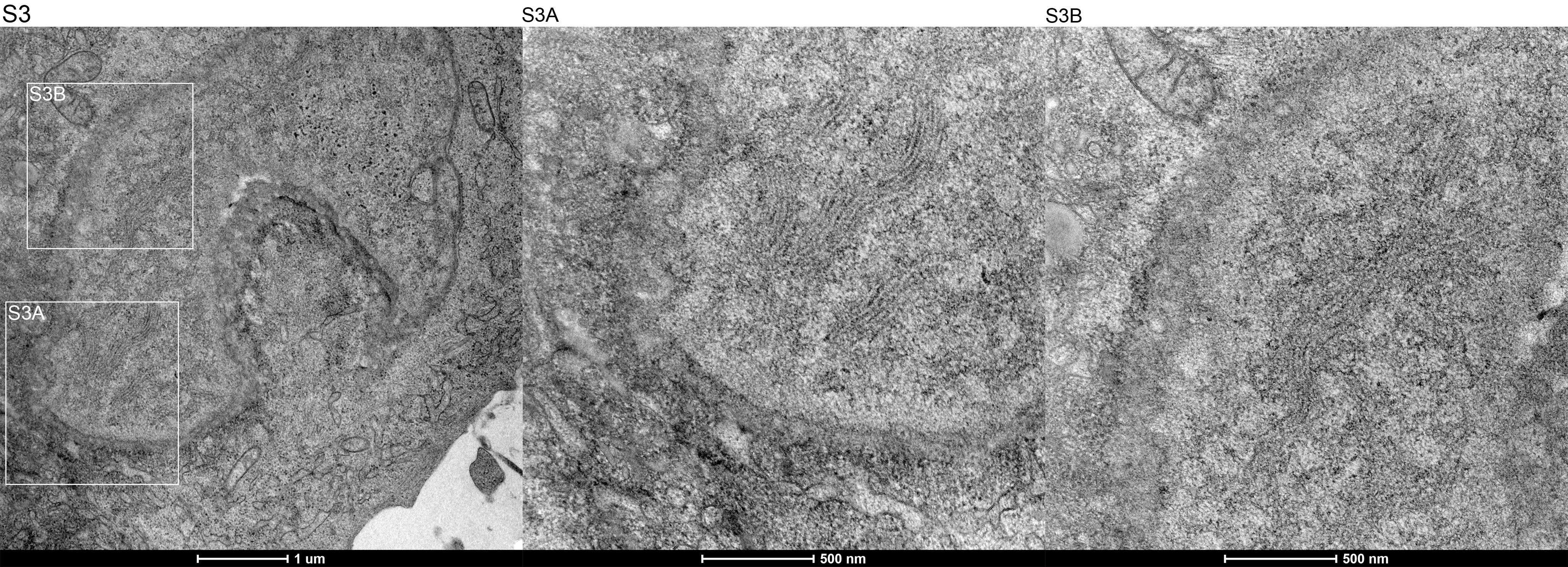
